# KinPred: A unified and sustainable approach for harnessing proteome-level human kinase-substrate predictions

**DOI:** 10.1101/2020.08.10.244426

**Authors:** Bingjie Xue, Benjamin Jordan, Saqib Rizvi, Kristen M. Naegle

## Abstract

Tyrosine and serine/threonine kinases are essential regulators of cell processes and are important targets for human therapies. Unfortunately, very little is known about specific kinase-substrate relationships, making it difficult to infer meaning from dysregulated phosphoproteomic datasets or for researchers to identify possible kinases that regulate specific or novel phosphorylation sites. The last two decades have seen an explosion in algorithms to extrapolate from what little is known into the larger unknown – predicting kinase relationships with site-specific substrates using a variety of approaches that include the sequence-specificity of kinase catalytic domains and various other factors, such as evolutionary relationships, coexpression, and protein-protein interaction networks. Unfortunately, a number of limitations prevent researchers from easily harnessing these resources, such as loss of resource accessibility, limited information in publishing that results in a poor mapping to a human reference, and not being updated to match the growth of the human phosphoproteome. Here, we propose a methodological framework for publishing predictions in a unified way, which entails ensuring predictions have been run on a current reference proteome, mapping the same substrates and kinases across resources to a common reference, filtering for the human phosphoproteome, and providing methods for updating the resource easily in the future. We applied this framework on three currently available resources, published in the last decade, which provide kinase-specific predictions in the human proteome. Using the unified datasets, we then explore the role of study bias, the emergent network properties of these predictive algorithms, and comparisons within and between predictive algorithms. The combination of the code for unification and analysis, as well as the unified predictions are available under the resource we named KinPred. We believe this resource will be useful for a wide range of applications and establishes best practices for long-term usability and sustainability for new and existing predictive algorithms.

## Introduction

Kinase regulation of protein phosphorylation plays important roles in almost all cell processes, including: growth factor/cytokine signaling, cell cycle and metabolic control, and DNA damage response. Serine/threonine phosphorylation-driven networks are anciently conserved and found throughout both prokaryotes and eukaryotes. Tyrosine phosphorylation is a more recent evolutionary mechanism, appearing as a complete signaling unit around the post-metazoan switch (1). We should also note that histidine phosphorylation plays essential roles in cell physiology, but experimental challenges for measuring it have hindered our ability to study and discover it (2), hence we focus on serine/threonine and tyrosine phosphorylation. These types of post-translational modifications (PTMs) are extensive in human biology and currently include roughly 40,000 tyrosine and 200,000 serine/threonine residues known to be phosphorylated (3). This number of discovered phosphorylation sites has grown dramatically in recent years (4), due to advances in mass spectrometry-based enrichment and discovery methods (5).

Unfortunately, our understanding of the role and regulation of the phosphoproteome has not kept pace with its discovery. For example, only about 3% of phosphorylation sites have a kinase that is known to recognize and phosphorylate that site (i.e. a kinase’s substrate), meaning that the vast majority of phosphorylation sites have no validated kinase (6). A major hurdle for validation is due to the preponderance of evidence required to ensure that a kinase is directly responsible for a specific phosphorylation site, especially when kinase networks often involve the activation of many kinases within a network. Since dysregulation of phosphorylation-driven networks are common to disease states and kinase inhibitors are a major class of FDA-approved drugs (7), it would be highly beneficial to connect dysregulated phosphorylation sites to candidate kinase targets.

A major breakthrough in the ability to hypothesize kinase-substrate connections came with the discovery that kinase catalytic domains have preferences for the protein sequence immediately surrounding the site of phosphorylation (8). However, kinases are often part of multi-domain protein architectures, where tertiary protein-protein interactions also help to shape kinase-substrate recognition (9, 10). Although full proteins may be referred to as substrates of kinases, it is individual tyrosine/serine/threonine residues that are phosphorylated by a kinase and not all sites on a protein are regulated by the same kinase. Hence, for the purposes of this work, the term *substrate* will refer to the site-specific phosphorylation site targeted by a kinase. The motivation to understand the kinases responsible for phosphorylating substrates and the discovery of kinase specificity determination has resulted in the proliferation of a large number of kinase-substrate prediction approaches, which extrapolate from available kinase-substrate data, or specificity features, to predict kinase-substrate relationships across the whole proteome. We found 31 such predictive algorithms published since 1999, ranging in methodologies from position specific scoring matrix approaches (11) to machine learning approaches, such as support vector machines (12, 13), Bayesian-based learning (14, 15), and recently, deep learning approaches (16). Additionally, new approaches that infer kinase networks from phosphoproteomic data have also recently emerged (17, 18) – although these types of predictions are on the order of the experimental size, versus at the scale of the entire proteome, which we focus on here. The myriad of approaches suggest a vast array of resources to consider when identifying candidate kinases for new phosphosites, integrating for network-based approaches, or looking for kinase enrichment, such as in KSEA (19).

Despite a plethora of resources, there are significant barriers to the use of these kinase-substrate prediction algorithms. For example, in our experience, more than one-third of the 16 resources published since 2010 are no longer accessible. In the event that the publication provided the predictions as supplementary material, there are two main classes of problems one encounters. First, there are issues with the sequences provided and mapping to current databases. For example, a phosphorylation site is often only designated by its position and an accession (e.g. Y1197 in UniProt record P00533). The older the dataset, the less likely there exists a Y/S/T at that position, as reference proteins are not static records. Second, these predicted resources might publish data filtered, either based on a stringency or based on the known phospho-proteome at that time. That presents a problem when one wishes to know whether the absence of a kinase-substrate edge represents a relationship that was tested and then filtered out due to stringency (a true negative), or is not present, because it was: 1) not tested (reference proteome difference) or 2) the site was not known to be phosphorylated at the time (reference phosphoproteome difference). Additionally, different age datasets and datasets published using alternate accession numbers are difficult to compare/cross-reference. The culmination of these challenges results in difficulty for researchers to consider multiple prediction algorithms in their research, to get complete results for the current scale of the phosphoproteome, or to interpret the meaning of various prediction algorithms.

To overcome these challenges, we built a process for unifying kinase-substrate prediction datasets, where substrates are residue-specific targets of kinases. This process uses a non-redundant reference proteome to retrieve proteome-level predictions (keeping all edges tested), maps kinase names to a common ontology, filters on the current phosphoproteome, and provides methods to allow for updates to both the reference proteome and phosphoproteome. Here, we present KinPred v1.0, a framework for unification, a unified resource containing three selected predictive algorithms, and analysis and methods for considering how to compare results, network sizes and emergent properties, agreement/disagreement between predictive algorithms, study bias, and the discriminability/overlap of different kinases within a predictive algorithm. We have made all code and datasets of KinPred v1.0 publicly available at https://github.com/NaegleLab/KinPred, and https://figshare.com/projects/KinPred_v1_0/86885, respectively.

## Results

### Evaluation of available kinase-substrate prediction sets

We performed a wide literature search of available kinase-substrate prediction algorithms. This encompasses approximately 30 prediction algorithms published between 1999 and 2020 (Table S1). Our goal was to select prediction resources that are: 1) available/accessible, 2) covered a large range of kinases, 3) residue-specific predictions, versus predictions of whole proteins as substrates, 4) proteome-scale predictions, and 5) kinase-specific, versus predictions by kinase family. We selected algorithms that focused on kinase-specific results, instead of kinase family predictions, such as DeepPhos (16), Quokka (20), and MusiteDeep (21), for the breadth of kinases and to allow comparisons between sets, but also because there is evidence that total sequence similarity does not guarantee similarity in specificity. Typically, the most recently duplicated paralogs may have high overall similarity, but the duplication event itself likely results in the freedom of one of these copies evolving new function – i.e. substrate targets (22). Instead, small changes in key regions that define specificity can occur (23), resulting in differential specificity, such as seen for the highly similar paralogs SRC and LCK (24). There are three prediction algorithms that met these requirements, which are: (NetworKIN (25, 26), PhosphoPICK (15), and GPS 5.0 (27) (referred to here as GPS). The three selected predictive algorithms use various features for prediction, which ultimately result in weighted edges between kinases and possible substrates, where the edge weight indicates a likelihood of that relationship, although edge weight values are not directly comparable across algorithms. NetworKIN’s (26) prediction approach uses a kinase-specific classifier, based on sequence and evolutionary phylogeny, and a network proximity score, based on annotated proximity between kinases and substrates from the STRING protein-protein interaction database (28). We used the combined score of the sequence-specific and network-specific scores (done by a naive Bayes approach), which is reported as a log-likelihood. GPS’s prediction approach relies on combined literature curation and kinase-substrate annotations from Phospho.ELM 7.0 (29) and PhosphoSitePlus (30), then uses a scoring approach based on sequence similarity to kinase substrates. GPS edge weights are reported as false positive rates based on generation of 10,000 random phosphopeptides, which are calculated within defined protein families based on protein kinase homology. Finally, PhosphoPICK, like NetworKIN, incorporates both sequencebased and network-based information. For training data, they used Phospho.ELM (29) and HPRD (31) for defined kinase-substrate relationships and BioGRID (32) and STRING (28) for protein-protein interactions. Their Bayesian network algorithm then estimates a likelihood p-value, which is the reported edge score. In summary, the three predictive algorithms rely heavily on sequence context, either from degenerate libraries (NetworKIN) or annotated kinase-substrate training sets (GPS, PhosphoPICK), where NetworKIN and PhosphoPICK add protein-protein interaction information and other context features.

### Integration of multiple kinase-substrate predictions into a common ontology that can be updated

Having selected our three predictive algorithms, we next wished to create a unified prediction set in which all possible human substrates had been tested and the substrates and kinases were mapped onto a common ontology. In this framework, there are two steps: 1) we used the algorithm to compute the prediction for all unique human proteins, capturing all predicted kinase-substrate edges – where edges may exist to Y, S, or T amino acids that are not known to be phosphorylated and 2) substrates were filtered to include only those sites that have been identified as phosphorylated according to the integrated compendia and experimental database of post-translational modifications, ProteomeScout, which combines five major compendia and primary phosphoproteomic experiments (3). The results are based on current ProteomeScout data (Feb. 7, 2020), which includes 41,414 unique phosphotyrosines and 204,399 unique phosphoserines/threonines. Combined, the three kinase-substrate predictive algorithms cover 76 tyrosine kinases and 266 serine/threonine kinases. Based on current UniProt annotations of human kinases this is out of 100 human tyrosine kinases and 376 human serine/threonine kinases possible. Of the few (9) dual specificity kinases, we include them in both categories, keeping their relevant substrates as either phosphotyrosine or phosphoserine/phosphothreonine substrates. Figure 1 shows the total kinases covered, indicating that GPS is the largest and PhosphoPICK is the smallest of the predictive algorithms in this set. This dataset guarantees coverage of known phosphorylation sites across the available kinases, all commonly mapped to the same kinase and substrate identifiers.

**Fig. 1.**
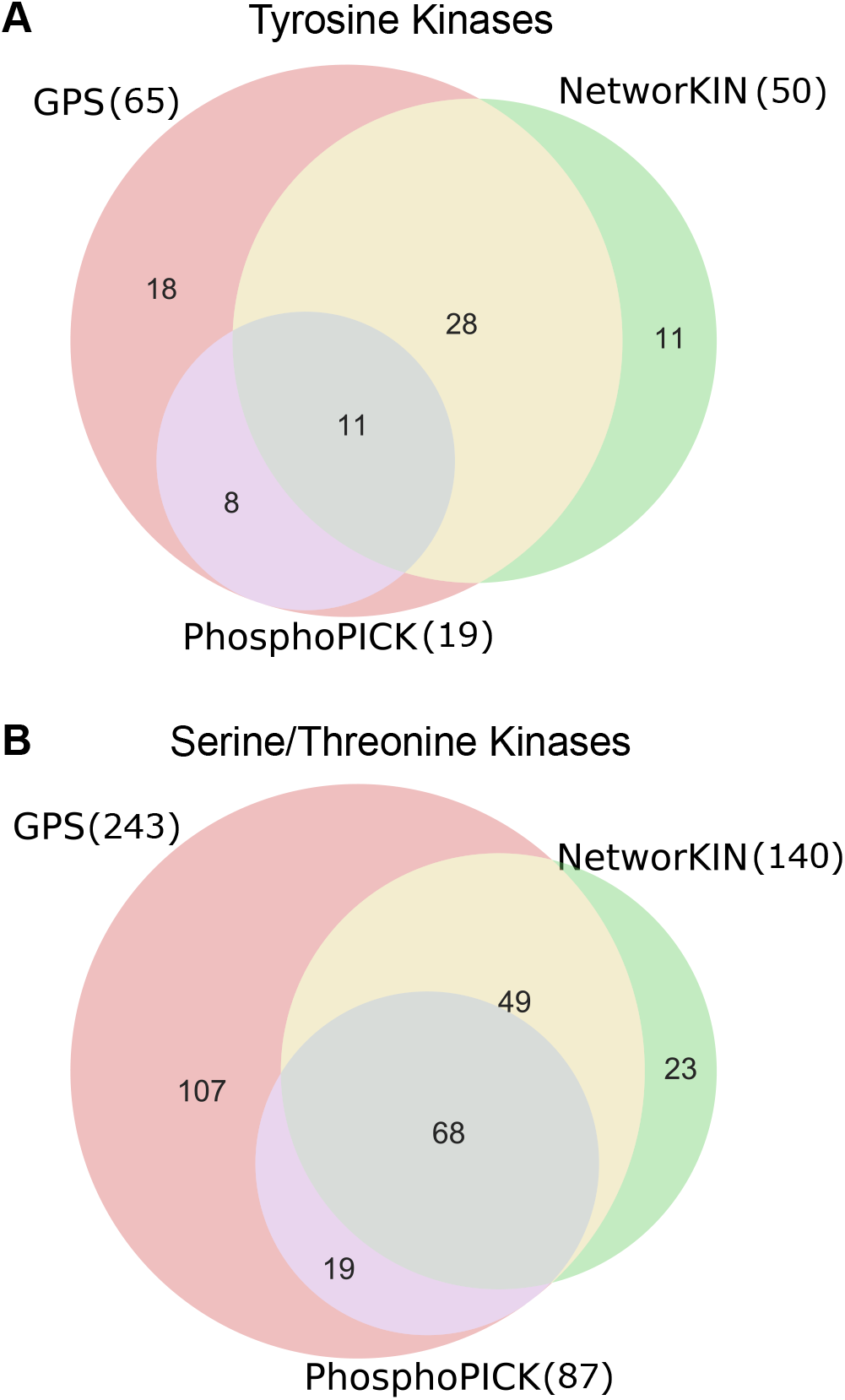
Comparison of the number of kinases in each predictive algorithm by kinase group. **(A)** Tyrosine kinases. **(B)** Serine/threonine kinases.

In order to update this common resource in the future when new phosphorylation sites are discovered, a trend that continues (4), we have implemented two basic approaches. First, most updates will require pulling pre-calculated edge weights from the resource that were filtered in the last version as not being phosphorylated at that time. Second, for a smaller number of cases, the underlying human proteome sequence may have changed in a way that affects the sequence surrounding the phosphorylated amino acid of interest. In these cases, our infrastructure identifies these protein sequences for running anew on the prediction servers/code, where new predictions are then replaced/added in the compendia of kinase-substrate predictions. We plan to update these predictions once to twice a year, when ProteomeScout undergoes a stable update. Hence, we provide here both a current, unified prediction set across the three predictive algorithms, but also a general framework for adding new predictive algorithms and updating prediction sets in the future.

### Thresholding kinase-substrate predictions

The kinase-substrate edges predicted by each algorithm have different mathematical interpretations and each resource publication suggested stringency thresholds (Table 1), where edges above that threshold are retained in the network. To explore the effects of these thresholds on overall algorithm coverage and size, we measured the number of unique phosphorylation sites in post-thresholded networks, across the suggested stringencies (Fig. 2A). There is a unique behavior between tyrosine and serine/threonine kinase networks – increasing stringency in tyrosine kinase predictions results in a significant loss of human phosphotyrosine coverage in NetworKIN and, to a lesser extent, PhosphoPICK. For PhosphoPICK, this may be related to the smaller number of kinases predicted, compared to the other algorithms. However, NetworKIN and GPS have more equivalent tyrosine kinases in their predictive algorithms (50 and 65, respectively). Hence, the decrease in NetworKIN phosphotyrosine coverage is likely due to the distribution of edges – fewer overall sites have at least one high stringency edge in NetworKIN, compared to GPS. This behavior is unique to phoshphotyrosine networks. Phospho-serine/threonine networks drop a significantly smaller fraction of the known human sites as a function of increasing stringency. Each algorithm stringency therefore has different effects on overall phosphorylation site coverage that is specific to the algorithm and the type of kinase network under consideration.

**Fig. 2.**
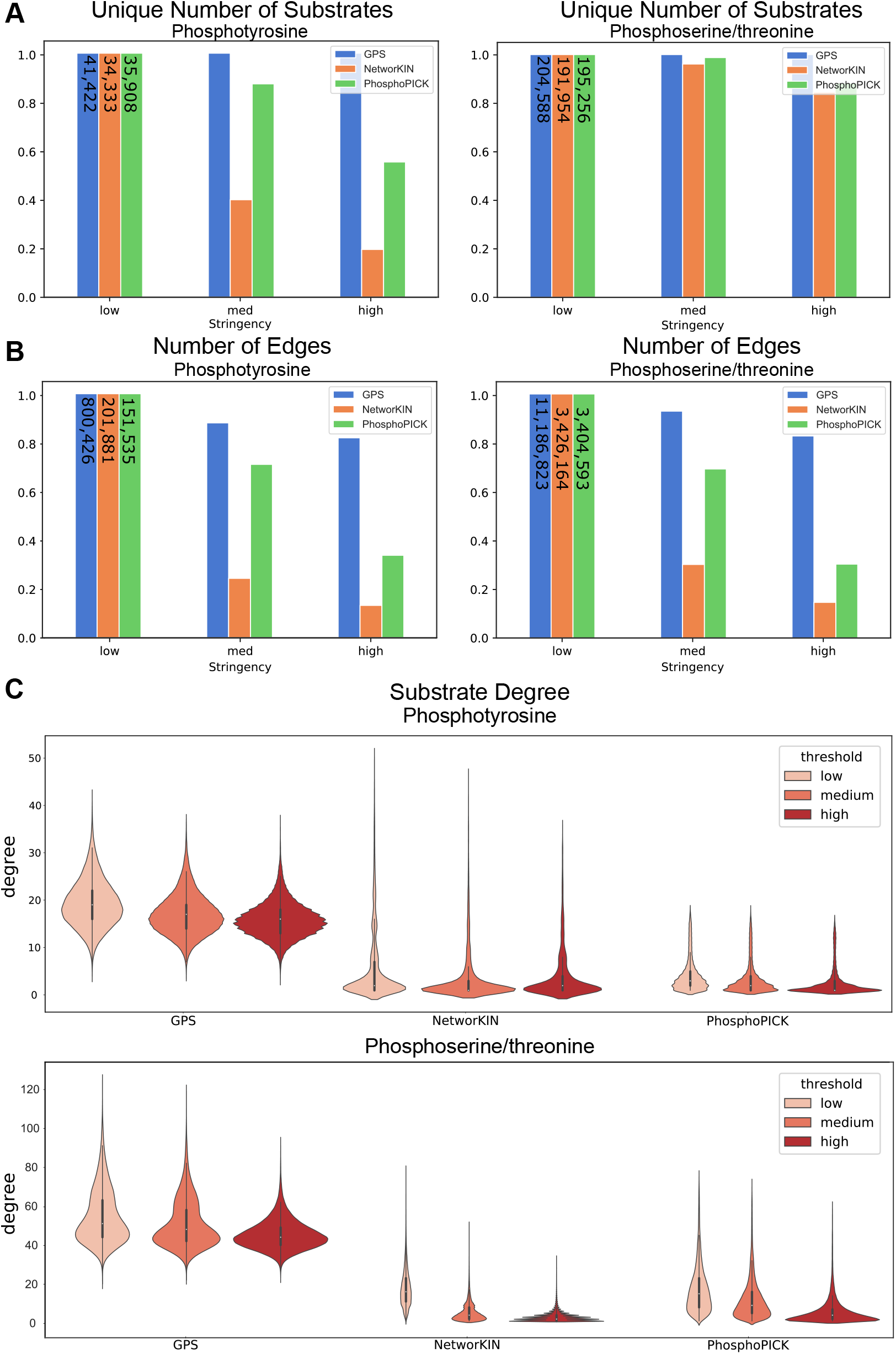
Network Comparisons Across Stringencies. **(A)** Bar graph of total phosphotyrosine and phoshposerine/phosphothreonine sites included in the network, normalized to the total number in the low stringency network, as a function of increasing stringency. Number of substrates at low stringency is labeled on each bar. **B)** Bar graph of total edges in the network, normalized by the total number covered in low stringency, as a function of threshold. Numbers on low stringency bars are the total number of edges at low stringency. **(C)** Distributions of substrate degree of phosphotyrosine and phosphoserine/phosphothreonine sites, where degree is the number of edges connected to a substrate at or above the denoted stringency.

**Table 1.**
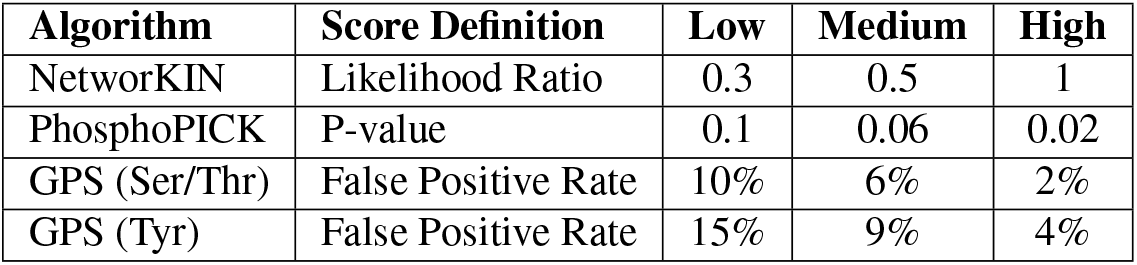
The definition of kinase-substrate edge weights in a given prediction network and stringencies cutoffs used by algorithm for low, medium, and high stringencies.

The total number of edges included in the post-thresholded networks is also of interest, this captures the density of kinase-substrate connections predicted in each network at, or above, the given stringency (Fig. 2B). At all thresholds, GPS contains the highest number of kinase-substrate edges, consistent with it being the largest of the three predictive algorithms, in terms of the total number of kinases available. However, the rate of loss of edges in the different predictive algorithms as a function of stringency differ. Specifically, medium and high stringencies in NetworKIN result in significant loss of predicted edges (as well as phosphorylation site coverage). To a lesser extent, this loss of network edges also occurs in PhosphoPICK.

Finally, in order to understand how edge loss translates to the connectedness of individual substrates, we explored the degree distribution of substrates as a function of stringency (Fig. 2C). Generally, the median substrate degree is directly proportional to the networks’ size (Fig. 2), where the center of the distribution is highest for GPS, which generally predicts significantly more kinases are responsible for phosphorylating substrates than the other algorithms. All algorithms demonstrate that there are a small proportion of substrates that are highly connected – being more than two standard deviations higher, compared to other substrates for that algorithm (Fig. 2C). For GPS, lowly connected substrates are considered outliers, which is different than the other algorithms, where the majority of substrates are lowly-connected – i.e. the majority of their substrates have connections from 0 to 10 kinase connections. Increasing stringency generally has the effect of reducing highly-connected outliers and decreasing the average degree of substrates. Taken together, the three different networks demonstrate very different properties regarding substrate inclusion and connectedness, including different effects of stringency thresholds, suggesting that direct stringency comparisons do not result in comparable networks.

### Effect of study bias in substrate prediction networks

The training of kinase-substrate predictive algorithms relies on well-annotated training data, which has the potential to create bias of strong predictions for phosphorylation sites that formed the basis of training sets, i.e. study bias. To test for this type of study bias, we looked at the relationship between the number of high-stringency edges a substrate has and how frequently it appears in compendia/databases of phosphorylation. ProteomeScout currently contains five compendia (PhosphoSitePlus (30), Phospho.ELM (29), dbPTM (33), HPRD (31), and UniProt (34)). Two of these resources, Phospho.ELM and HPRD are no longer updated, but represent the oldest data that was the foundation of many kinase-substrate prediction algorithms. Conversely, PhosphoSitePlus, the largest resource, is predominantly composed of mass spectrometry-based discovery and accounts for a large proportion of the data represented by just one compendium. Interestingly, none of the compendium completely overlaps with another compendium. Hence, we propose that the total number of compendia a phosphorylation site appears in can be used as a proxy for how likely they were included as part of the original training or testing data of kinase-substrate prediction algorithms – i.e. a measure of training set study bias.

Figure 3 shows the distribution of substrate degrees as a function of the number of compendia the substrate has been observed in. Roughly 93% and 90% of phosphotyrosine and phosphoserine/threonine sites, respectively, are annotated by at least one compendium, the majority of which are annotated only in one compendia (60% and 53% respectively). Roughly 16% of phosphotyrosines and 23% of phosphoserine/threonine are very well studied, appearing in at least three different compendia. Most predictive algorithms, in both phosphorylation groups, generally show increasing degree with increasing compendia annotations, suggesting a strong relationship between how many high-stringency edges a substrate has and how well-studied it is. To statistically test this, we used the Kolmogorov-Smirnov (KS)-test to compare the distributions of a group to the distribution in the next compendia group. If substrates were truly a random sampling of the background, the KS-test would result in no significant rejections of the null hypothesis. However, we observed very significant differences in most comparisons, including all of the tests in GPS substrate degrees and most of PhosphoPICK tests. NetworKIN demonstrates fewer significant differences in network degree as a function of study bias. These results suggest that it is important to understand the effect of study bias that underlies the kinase-substrate predictions when drawing conclusions from these predictive algorithms.

**Fig. 3.**
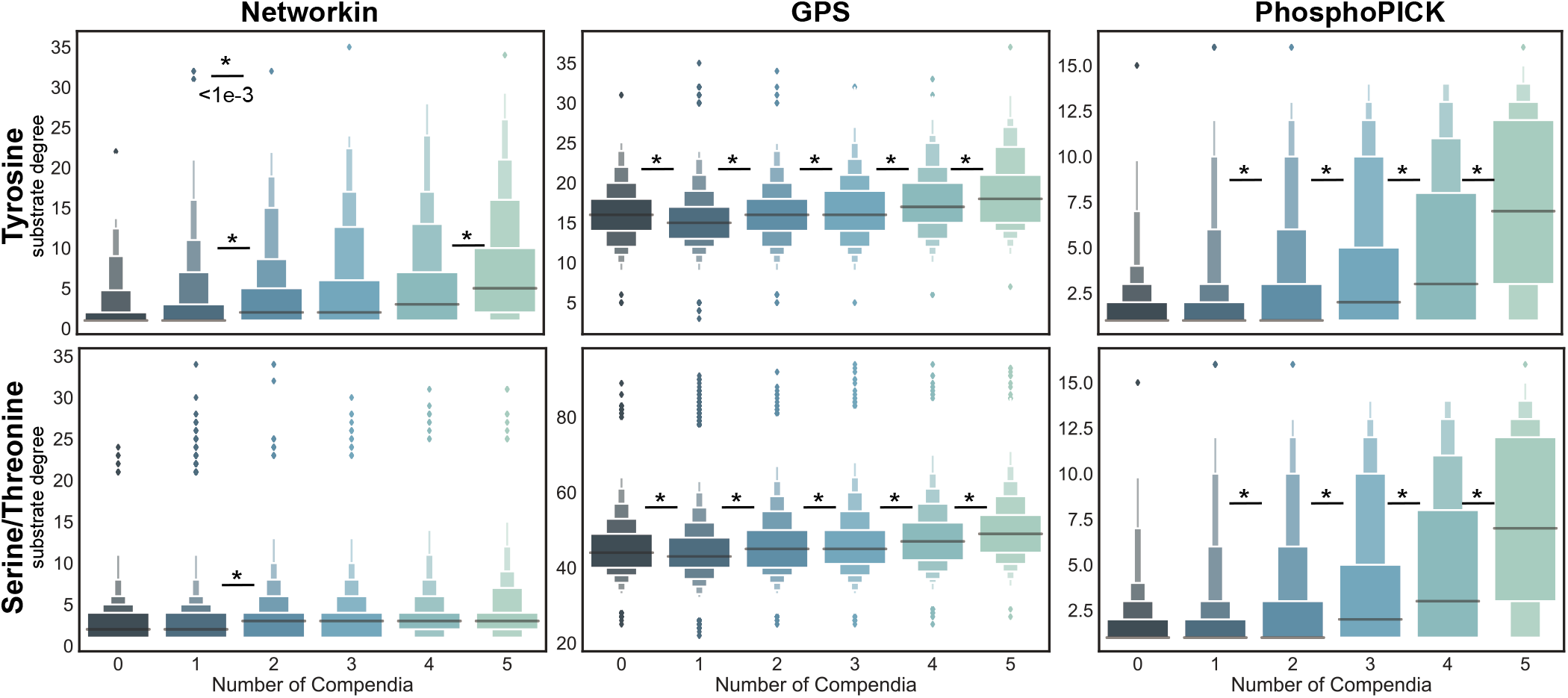
Effect of study bias. Each panel shows the distribution (boxen plot) of the number of high stringency kinase connections a substrate has, where substrates are grouped according to the number of compendia they were annotated in, where the more the compendia a substrate is in the more likely it was part of a training set for the algorithms (i.e. a type of study bias). Distributions are plotted for high-stringency networks for NetworKIN, GPS, and PhosphoPICK (from left to right) and for tyrosine (top) and serine/threonine networks (bottom). To test if there is a difference between one compendia group and the next higher compendia group, we performed pairwise Kolmogorov-Smirnov (KS)-tests, where * denotes <1e-3 p-value, indicating the data do not come from the same underlying distribution, which is the expected behavior if each sample distribution by compendia was randomly pulled from the background.

### Direct comparison of kinase-substrate predictive algorithms

We wished to understand the similarity of kinase-substrate predictive algorithms. For these analyses, we consider only the kinases that are predicted in any pair of predictive algorithms (Fig. 1), which includes: 11 tyrosine kinases and 68 serine/threonine kinases between PhosphoPICK and NetworKIN; 39 tyrosine kinases and 117 serine/threonine kinases between GPS and NetworKIN; 19 tyrosine kinases and 87 serine/threonine kinases between GPS and PhosphoPICK. One aspect of comparison we considered was whether different algorithms generally agreed with how connected kinases are – i.e. their degree or number of substrates predicted per kinase. To measure this, we looked at the correlation in kinase degree between predictive algorithms at different stringencies. There was no discernible correlation between any pair of predictive algorithms. The highest correlation was between PhosphoPICK and NetworKIN tyrosine kinase degrees with a Pearson correlation of 0.49 (p-value of 0.12, likely due to few numbers of kinases in the set). If we perform the same analysis from the perspective of substrate connectivity, there is a modest correlation only between PhosphoPICK and NetworKIN for both tyrosine (r=0.6, p-value < 1e-100) and serine/threonine networks (r=0.4, p-value < 1e-100). All other pairwise comparisons of kinase and substrate degree showed poor correlation. In short, this means algorithms do not generally agree about which kinases and substrates are highly or lowly connected within networks and only PhosphoPICK and NetworKIN have some agreement as to the general trend of substrate connectivity.

Finally, we wished to ask how similar/dissimilar kinases are within and across predictive algorithms. We measured the similarity between kinases using the Jaccard index, which is the size of the intersection of two sets, normalized by the size of the union of the sets, for thresholded networks. The Jaccard index ranges from 0 (no overlap), to 1 (complete overlap). There is an expectation that some amount of overlap occurs by random chance. For sets containing at least 100 elements, a Jaccard index of 0.49 or larger is considered significant at a p-value of 0.001 (35). A kinase set is defined as the substrates connected to it at, or above, a given threshold. Since all comparisons here contain at least 100 elements, then 0.49 is a bound with conservative estimate of being unlikely due to random chance alone. Additionally, a Jaccard index of 0.49 corresponds to roughly half of the available substrates between a pair of kinases as being shared. We used this approach to understand network overlap between kinases within the same algorithm and between kinases in different predictive algorithms.

#### Within-algorithm kinase similarity

We compared within-algorithm kinase similarity to understand the overlap, or lack of discriminability, between kinases within an algorithm. Figure 4 shows the results of all pairwise Jaccard scores for tyrosine kinase predictions within NetworKIN at low and high stringency. The highest degree of overlap exists at low stringencies, where 46% (23) of the tyrosine kinase networks predicted by NetworKIN are significantly similar to at least one other tyrosine kinase in NetworKIN. This overlap decreases with increasing stringency – dropping to 26% of kinases (13) at medium stringency and 22% of kinases (11) at high stringency. The NetworKIN serine/threonine kinase overlap starts higher (65.7% overlap significantly at low stringency), but also drops faster to end at only 9.3% overlap at high stringency. PhosphoPICK shows no significant overlap at high stringency for tyrosine kinases and modest overlap (2%) for serine/threonine kinases. The general trend of decreasing overlap with increasing stringency is consistent across all predictive algorithms and both tyrosine and serine/threonine networks (Fig. S1), with the exception of tyrosine kinase networks in GPS, which actually increase in the extent of overlap with increasing stringency.

**Fig. 4.**
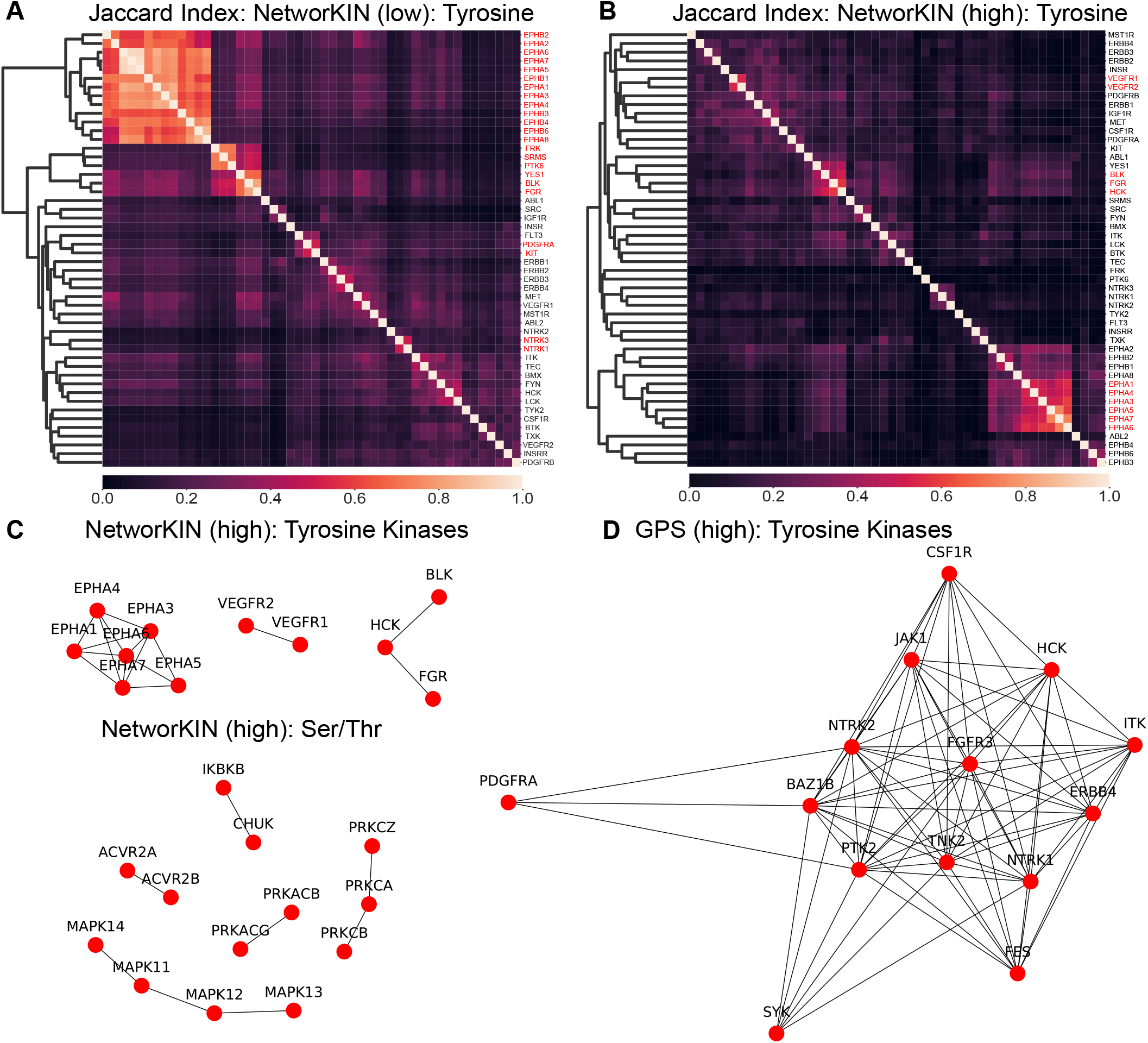
Within-algorithm kinase similarity. Jaccard index matrices for NetworKIN predictions at **(A)** low and **(B)** high stringency. The symmetric matrix was sorted hierarchically and kinase names are highlighted in red text if their Jaccard index meets a minimum of 0.001 p-value significance (a value of 0.49 or greater). Those relationships that exceed significant Jaccard index values are shown in network diagrams **(C)** NetworKIN tyrosine and serine/threonine kinases under high stringency and **(D)** GPS tyrosine kinase networks (serine/threonine are large and available in Fig. S1). A spring layout was used on the weighted graph, so the closer the nodes the larger the Jaccard index. These graphs show all significant overlapping kinases in the given kinase network.

We next used a network layout to see what kinases significantly overlap with each other (Fig. 4C,D and Fig. S1). The kinases that significantly overlap with each other show very different patterns in NetworKIN than in GPS. In NetworKIN high stringency networks, overlap occurs between highly homologous paralogs of kinases (e.g. the EPHA and MAPK family of kinases). However, in GPS there is low overlap amongst these families, but high overlap between evolutionary diverse kinases, such as CSF1R, JAK1, ITK, ERBB4, and PTK2. Hence, our analysis suggests that when using the kinase-substrate networks as a whole, one must consider the degree of overlap, i.e. lack of discriminability, between kinases in an algorithm and the role of stringency in alleviating, or in the case of GPS, increasing overlap.

#### Between-algorithm kinase similarity

We next wished to understand whether the different predictive algorithms generally agreed regarding kinase network similarity. We again used the Jaccard index, this time measuring overlap between a kinase in one algorithm and kinases in a second algorithm. Most notably, this analysis demonstrated that globally the between-algorithm similarity of the same kinases is significantly lower than within-algorithm similarities of different kinases. For example, Figure 5 shows NetworKIN vs. Phospho-PICK at low stringencies for both tyrosine and serine/threonine networks (all direct comparison JI matrices are available in Fig. S2), with JI values that are significantly lower than the best JI values seen within an algorithm (Fig. 4A).

**Fig. 5.**
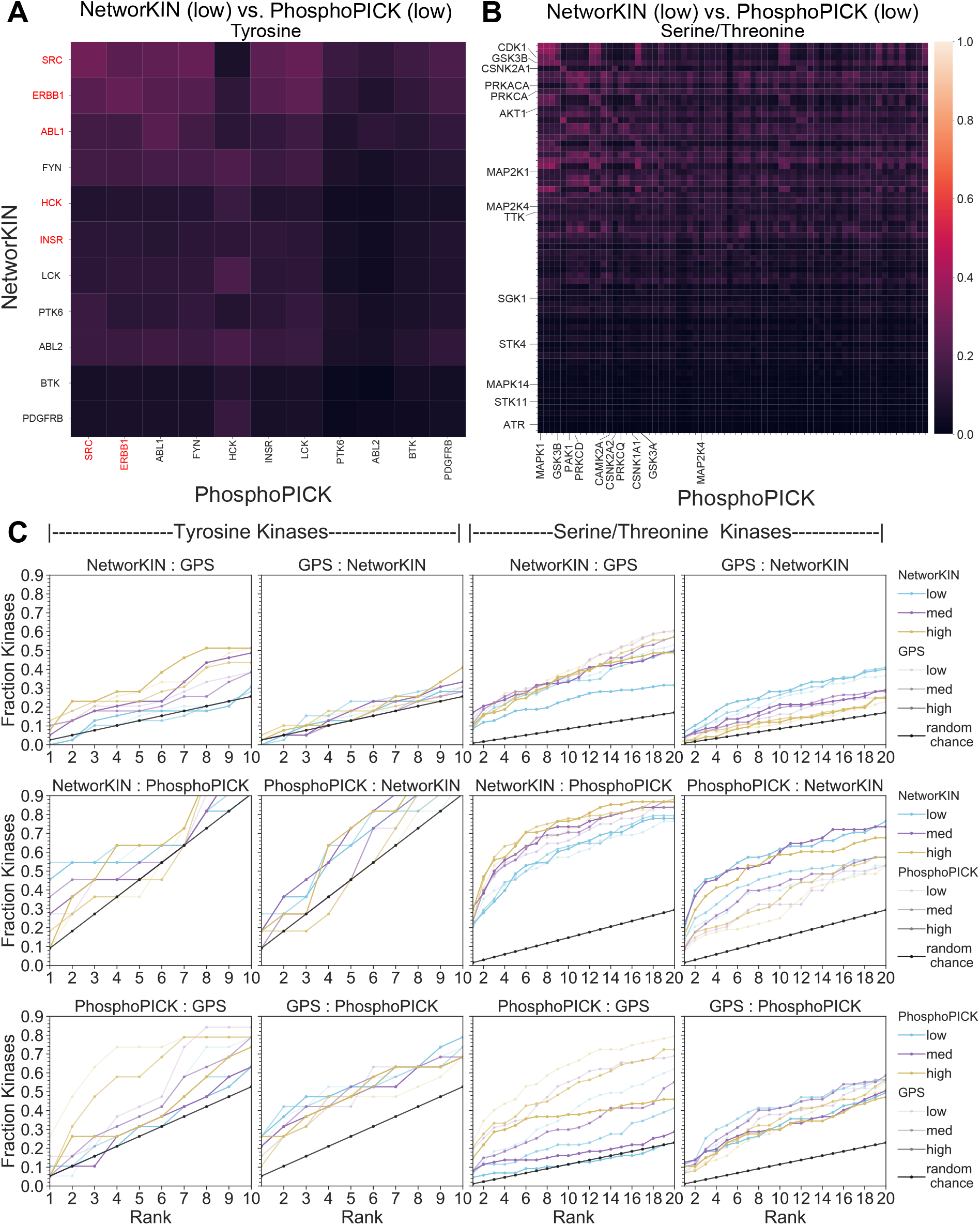
Between-algorithm kinase similarity. Jaccard index matrices (non-symmetric), comparing GPS and NetworKIN, both at low stringency, where highlighted labels in red for tyrosine kinases **(A)** or labeled for serine/threonine kinases **(B)** indicate that comparison falls within the top 1% of all kinase comparisons. **(C)** In order to understand if the highest overlap of a kinase in one algorithm is the same kinase in the other algorithm, we calculated the cumulative distributions of the fraction of this match from the highest ranked kinase to the top 10 (tyrosine kinases) or top 20 (serine/threonine kinases). For example, a y-axis value of 0.3 at x-axis value of 2 means that 30% of the kinase comparisons from algorithm A to algorithm B, where the title defines A:B comparison in that panel, have a JI value that puts them in the top 2 of all kinase-kinase comparisons. The black line gives an estimation of what is expected by random chance. This calculation was run for all possible thresholds and the color line is used to indicate the threshold of algorithm “A” and the thickness indicates the threshold of the “B” algorithm. For example, the top-left panel, the thick gold line indicates the comparison of NetworKIN’s high-stringency network to GPS high-stringency network, which also has the largest deviation from the random-chance line of all possible NetworKIN:GPS comparisons. The faster and higher the deviation from the random chance line, the more agreement there is between prediction algorithms networks at those stringencies.

Despite having low overall similarity, we wished to know if the best overlap still occurred between predictive algorithms for the same kinase. Figure 5 and Figure S2 highlight the kinases whose match is in the top 1% of Jaccard scores measured against all kinases in another algorithm. We noticed that although some kinase’s rank high when comparing from one algorithm to another, the opposite is not necessarily true. For example, SRC, ERBB1, ABL1, HCK, and INSR rank high when comparing NetworKIN to PhosphoPICK, but only SRC and ERBB1 rank high when comparing from PhosphoPICK to NetworKIN. Based on Jaccard scores, HCK of NetworKIN overlaps best with HCK PhosphoPICK networks, but the HCK PhosphoPICK network instead overlaps to a greater extent with NetworKIN’s LCK, ABL2, and FYN networks. The same trend holds true for serine/threonine networks, where 14 kinases in NetworKIN’s overlap in the top 1% of rankings with PhosphoPICK, but only 10 PhosphoPICK kinases overlap in the top 1% of JI values with NetworKIN kinases. Finally, we noted that the identity of the best-ranking kinases is highly dependent on stringency – the kinases that overlap change drastically across different stringency networks (Fig. S2).

Since each algorithm’s stringency value has a different interpretation and results in different impacts on the overall networks (e.g. the number of edges, edge degree distributions, etc.), we wished to compare all combinations of stringency in a pairwise manner to identify the maximum overlap of kinase networks between predictive algorithms. Figure 5 shows the results of this experiment, presented as the cumulative distribution function of the kinases by rank – meaning one can measure the fraction of kinases whose direct match appears in the top 10 or top 20 ranks for tyrosine and serine/threonine networks, respectively. We included a line (in black) that establishes a bound on random expectation for this type of calculation. Additionally, we randomized the networks and performed this experiment, which demonstrated that random networks generally follow random expectation (Fig. S3).

The CDF ranks, akin to the findings of the top 1% of kinases in the heatmaps, highlights that the asymmetry in mapping results is universal. Specifically, there are drastically different behaviors in mapping from algorithm “A” to algorithm “B” than from B to A. For example, not only are the starting points (fraction that are ranked first) in the A:B than B:A plots different, but which stringency selections achieve optimum overlap varies between comparisons and between kinase types. For example, low stringency NetworKIN predictions have the best overlap with PhosphoPICK for tyrosine kinases, but the opposite is true for the serine/threonine comparison, where high stringency creates the best overlap. Additionally, the shape of the curve, where positive deviation from the random line indicates more similarity between predictive algorithms, is not the same in the A to B than the B to A comparisons. For example, although mapping from NetworKIN into GPS deviates positively from the random line, the opposite is not true – GPS rankings are not better than expected by random chance. This means that GPS overlap of kinases with the same kinase in NetworKIN is indistinguishable from the overlap they share with different kinases in NetworKIN. GPS mappings in both kinase groups and against both NetworKIN and PhosphoPICK perform the worst in matching relevant kinases in other predictive algorithms. Taken together, these results suggest that there is generally low agreement between predictive algorithms, although it is often better than random chance, suggesting at least some agreement. Unfortunately, these results also suggest that there is no single best stringency to recommend that maximizes the comparisons between predictive algorithms.

## Discussion

### Lessons from comparisons of multiple predictive algorithms

The analyses we performed on the three different prediction datasets demonstrated vastly different behaviors in terms of network size, properties, study bias, and overlap both within and between predictive algorithms. Sadly, the deeply disparate results from the network overlap analysis suggest that it is unhelpful to merge networks in a manner that relies on agreement between predictive algorithms. Instead, users of these networks should likely consider specific needs when determining which predictive algorithms to use, how to consider stringency, and how to aggregate results between predictive algorithms. For example, if one wishes to minimize overlap between kinase networks within an algorithm, then for serine/threonine predictions you would select high stringency networks for all predictive algorithms, but for tyrosine networks you would select low stringency for GPS, medium or high stringency for PhosphoPICK, and high stringency for NetworKIN. If instead, one wished to maximize network coverage of phosphorylation sites, then GPS at any stringency and low stringencies for NetworKIN and PhosphoPICK would be ideal. Finally, if the goal is to maximize overlap between predictive algorithms, there is no single stringency selection that is ideal, even in a single pair of predictive algorithms, since mapping from one to the other is not equivalent in the reverse mapping. In conclusion, we propose that the general analysis framework developed here will be useful to researchers when considering this, or other predictive algorithms.

Another important conclusion from this analysis, is that it is likely that the predominate behaviors within the predictive algorithms are largely driven by the underlying assumptions and training data at the basis of the predictive algorithms. For example, NetworKIN’s basis for sequence recognition did not come from annotated sets, but from degenerate libraries and therefore, may be the reason that NetworKIN does not exhibit the same extent of study bias within its network as GPS and PhosphoPICK. Since the latter two predictive algorithms relied on relatively small numbers of kinase-substrate annotations coming from databases like Phospho.ELM, we hypothesize that these predictive algorithms enrich for confident scores on well-studied proteins from the training data, which results in the very high degree of study bias observed. Secondly, NetworKIN’s poor discrimination between highly homologous kinases is likely due to its reliance on assumptions that homologous kinases result in similar specificity. Unfortunately, whether highly homologous kinases are more likely to phosphorylate similar substrates, versus the suggestion of GPS, where highly dissimilar kinases have high degree of overlap, is difficult to explore given the small amount of available, validated kinase-substrate relationships. Beyond the onus of explicit kinase-substrate identification, recent approaches for assessing kinase specificity profiles have been developed, which demonstrated that two highly homologous proteins (SRC and LCK) have evolved different specificities (24). Ideally the expansion of these approaches will eventually aid in the determination of the relevance of homology-based specificity assumptions in kinase models.

### Value of cohesion and growth

KinPred (version 1.0) is the first bioinformatics resource that provides a unified kinase-substrate prediction network for the human phospho-proteome from multiple predictive algorithms. Here, we retrieved unified results for all kinase-specific predictive algorithms that were accessible at the time of this research. We believe this is a useful resource for any researcher wishing to explore specific kinase-substrate hypotheses or for use in algorithms that rely on entire networks, such as KSEA (19). In addition to providing current, updated results of these three predictive algorithms for the whole proteome, we have provided code and a created a general framework for: 1) unifying new approaches in kinase-substrate predictions 2) updating resources as knowledge of the human phosphoproteome grows, and 3) updating resources for when the underlying protein databases are updated in a way that impacts specific kinase-substrate predictions. By using a framework for updating these predictions, it helps to ensure the usability, accessibility, and accurateness of these resources for future users.

Unfortunately, our broad exploration of kinase-substrate prediction resources highlighted challenges users face in accessing or understanding the predictions. Of the five predictive algorithms for site-specific kinase-substrate prediction, three were unavailable or unaccessible for creating updated results. For available resources, if we wished to guarantee coverage of the human phosphoproteome, it required running the full proteome against the website, or in downloadable code. This process took several weeks for the largest predictive algorithms. Hence, there are significant barriers to the usability of these resources and we would like to emphasize that for long term sustainability/usability of new algorithms in the field, publication should include the entirety of the prediction against a reference proteome (i.e. all kinase-substrate edges tested) and a subset of those predictions for the current phosphoproteome, provided with the reference proteome and phosphoproteome used. This way, should algorithm access no longer be available, users could more easily access and update from the larger set of available predictions provided at the time of publication. Ideally, if prediction results were provided in the unified format we have developed, results are also easily comparable between algorithms. Finally, an advantage to providing all edge weights, instead of filtering on a specific stringency, is that, as we demonstrated, stringency selection for considering edges as significant depends on the application, so publishing only post-filtered results presents a significant barrier for the use of these resources.

## Materials and Methods

### Data and Code Availability

All retrieval, processing, and analysis code was written in Python3/Jupyter and are available in our Github repository at: https://github.com/NaegleLab/KinPred. All datasets, midpoints, reference proteome and phosphoproteomes, and mapping ontologies, extracted code, for KinPred v1.0 are available on Figshare at: https://figshare.com/projects/KinPred_v1_0/86885.

### Data Preparation

#### Retrieving and processing a unified prediction set

We used a non-redundant canonical human proteome reference file from UniProt (Feb. 26, 2020, stored with project data at: https://doi.org/10.6084/m9.figshare.12749342.v1). The downloaded FASTA file was programmatically split into 21 smaller files and then submitted to the prediction services, using the appropriate programmatic or web-based interface, as outlined below. We removed prediction types that did not refer to protein kinase-substrate interactions. For unification between predictive algorithms, the raw prediction data was pre-processed to a standard format, and a controlled vocabulary was used (labeled Whole proteome-level predictions in the Figshare project). The substrates and kinases were mapped to the same ontology, compliant with the UniProt reference. The standardized format includes the following information: the unique identifier for substrates, which is the combination of the protein UniProt accession and site position within the protein sequence; the protein accessions and gene names annotated in UniProt; the site, which is the combination of the amino acid residue and the site position within the protein sequence; the peptide sequence around the predicted site; and the common kinase name used across all predictions. Each algorithm has a pre-processed prediction data file with a standardized format. We created a global kinase ontology reference to map between algorithm-specific kinase names and a common kinase name (available at: https://doi.org/10.6084/m9.figshare.12749333.v1). The global kinase map contains the UniProt accession, a common kinase name, the description of the kinases, the kinase type, and the reference kinase names in each algorithm. Additionally, this file contains a preferred name, so that users may define preferred kinase names in downstream analyses.

##### NetworKIN

predictions were run from a local installation of NetworKIN3.0, software available at https://networkin.info/download/NetworKIN3.0_release.zip. Predictions were set to include all kinases with no threshold, in order to retrieve all possible edges, and Human was the selected species, if available.

##### GPS

The GPS5.0 application was downloaded (GPS5.0 Unix v. 20190721 from http://gps.biocuckoo.cn/download.php), installed locally, and predictions were initially collected in January 2020, then updated to Feb. 26, 2020 reference proteome.

##### PhosphoPICK

predictions were retrieved in November 2019 by submitting the split FASTA files to the web interface at http://bioinf.scmb.uq.edu.au/phosphopick/submit. Initial predictions were performed on a reference proteome from Oct.16, 2019, and then updated to the Feb. 26, 2020 reference proteome.

### Filtering predictions for known phosphorylation sites

We retained kinase-substrate predictions for known phosphorylation sites by using the reference phosphoproteome from ProteomeScout (downloaded Feb. 2020, reference available in project data with the human UniProt reference on the Figshare project in the Raw directory). Phosphosites were retrieved from the ProteomeScout database by protein accessions using ProteomeScout API (4, 36) for every UniProt accession in the reference proteome. The phosphorylation sites were matched by the site position. If the site position did not match between the ProteomeScout UniProt record and the reference record, we matched the substrate sites by using the algorithm returned 11-mer or 15-mer (GPS, PhosphoPICK) peptide – which, for some highly repetitive sequences in the human proteome, may match multiple times in a single protein record. In these cases, all matches were retained in the final data format. Kinase-substrate predictions were filtered for coherence – i.e. serine/threonine sites were removed if the kinase is a known tyrosine kinase and vice versa. Tyrosine/serine/threonine residues were kept for dual specificity kinases only. The final datasets for each algorithm are available in the Figshare project under the “Final Data” directory. It is important to note that no stringency filters have been applied to this data, so that researchers have full control over edge weights of interest. However, all analyses in this paper applied at a minimum the lowest stringency as defined in Table 1.

#### Updating prediction data

We wrote methods to update the human phosphoproteome predictions for future releases of ProteomeScout datasets. This method checks for changes between the human reference used to generate KinPred v1.0 and a new human reference dataset. Any sequences that have changed or been added will be provided, such that the retrieval of new predictions can be performed and added/replaced in the datasets. Next, phosphosites in the new reference phosphoproteome, mapped to the common substrate ontology, are used to filter the whole proteome-level predictions for each resource.

### Data Analysis Methods

#### Thresholds for Final Prediction Data Comparison across predictive algorithms

Each algorithm has a different interpretation of what an edge weight signifies and therefore, there is no single value that can be applied to threshold all predictive algorithms. The thresholds used for filtering each algorithm dataset are summarized in Table 1.

##### NetworKIN

the unified likelihood was used as the score of the predictions. The unified likelihood of a certain NetPhorest probability and network proximity score of STRING were calculated by the joint probability rule (26). Horn et al. indicated that it is likely to be true for the unified likelihood scores to be higher than the theoretically neutral value of 1 (26). Therefore, for the purpose of the algorithm comparisons for NetworKIN prediction data, the low, medium, and high thresholds were adopted with the unified likelihood of 0.3, 0.5, and 1, respectively.

##### GPS5.0

the default low, medium, and high thresholds for serine/threonine kinases were adopted with FPRs of 10%, 6%, and 2%, respectively; and 15%, 9%, and 4%, respectively, for tyrosine kinases (27). These default thresholds were kept in this study. To obtain the actual cutoff scores for the predicted kinases, a random subset of the human proteome sequences was submitted to GPS5.0 with threshold setting of low, medium, or high. The actual cutoff scores were extracted from the ‘Cutoff’ column of the prediction results of GPS5.0 application. These values are provided in the Figshare project under the Raw/GPS directory.

##### PhosphoPICK

PhosphoPICK is the only resource that includes a fourth category of stringency, with default thresholds of 0.25, 0.1, 0.05, and 0.005 as the low, medium, high, and very high, respectively (15). In order to identify a relevant set of cutoffs that produce three categories for direct comparison to GPS and NetworKIN, and since both GPS and PhosphoPICK stringencies are based on a false positive rate, we used p-values of 0.1, 0.06, and 0.02, for low, medium, and high stringencies, respectively.

#### Predicted Kinase Networks Similarity Comparisons

The predicted kinase network similarity comparisons were measured by the Jaccard’s Index (35). The Jaccard’s index is the ratio of the intersection to the union of the two sets. The values range from 0 for no similarity to 1 for the complete overlap of the two compared sets. Based on “Tables of significant values of Jaccard’s index of similarity” (35), the lower and upper critical values of Jaccard’s index with the probability levels 0.05, 0.01 and 0.001 were calculated for the N elements in either of the two compared OTUs (Operational Taxonomic Units). The upper critical values of Jaccard’s index of 0.49 with N=100, which is the upper limit for the calculated table of the significant values of Jaccard’s index, correspond with the probability level of 0.001. Hence, we used a threshold of 0.49 to identify similarities between kinases that are significant.

## ACKNOWLEDGEMENTS

Research reported in this publication was supported by the National Cancer Institute of the National Institutes of Health under Award Number R21CA231853. The content is solely the responsibility of the authors and does not necessarily represent the official views of the National Institutes of Health.

---

**Fig. S1. Within Algorithm Similarities** Within-algorithm kinase similarity measured by Jaccard index for all predictive algorithms at all stringencies (low, medium, and high from left to right). The symmetric matrix was sorted hierarchically. The within-algorithm relationships that exceed significant Jaccard index (0.49) values are shown in network diagrams, below their corresponding heatmap. The figures are separated by kinase type (tyrosine kinases first and serine/threonine kinases second. For dense maps, where it is not possible to achieve label separation, the names of kinases can be found as labels in the heatmap, where heatmap sorting correlates with proximity in the network graphs.

**Fig. S2. Between Algorithm Similarities** Between-algorithm kinase similarity measured by Jaccard index for all combinations of predictive algorithms at all stringencies (low, medium, and high from left to right). The matrices are sorted by the same kinase order on both the Y- and X-axes. If the ranking of kinase in algorithm A is in the top 1% of similarities in algorithm B comparisons, the font is in red for tyrosine kinases, or indicated with a label in serine/threonine kinases.

**Fig. S3. Ranking Performance for Randomized Datasets** The edges of predictive algorithms were randomized and the process of measuring Jaccard index overlap between predictive algorithms and measuring the cumulative distribution function of rankings, as done in Figure 5 was repeated on random datasets. The dashed-black line indicates the expectation of random performance line.

**Table S1. Overview of predictive algorithms**. The name, Pubmed link, and details about the availability and details of 49 predictive algorithms of phosphorylation and kinase-substrate relationships.

**Table S2. Ranks of kinases in between-algorithm comparisons** This Excel book contains an overview of the performance of each pairwise algorithm overlap, the optimum stringency selected, and the ranks of kinases mapping from one algorithm to another at those optimum stringencies.

## Notes

https://github.com/NaegleLab/KinPred

https://figshare.com/projects/KinPred_v1_0/86885

### Competing Interest Statement

The authors have declared no competing interest.

### Summary of Updates

Minor revisions of text to increase clarity.

