## Supplemental Figures for "KinPred: A unified and sustainable approach for harnessing proteome-level human kinase-substrate predictions"

#### Supplemental Figure 1: Within Predictor Similarities

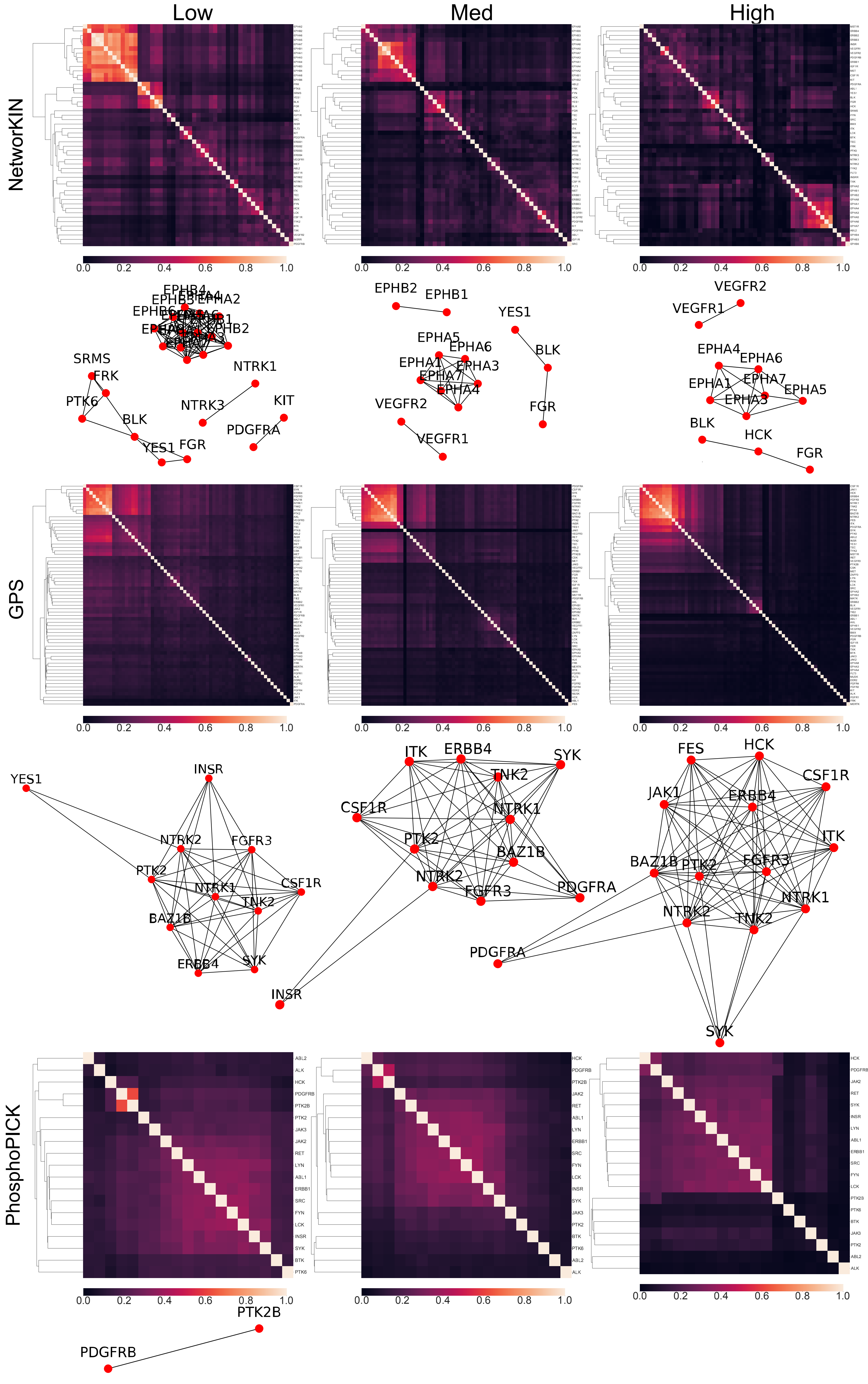

### Supplemental Figure 1: Within Predictor Similarities, continued (ser/thr)

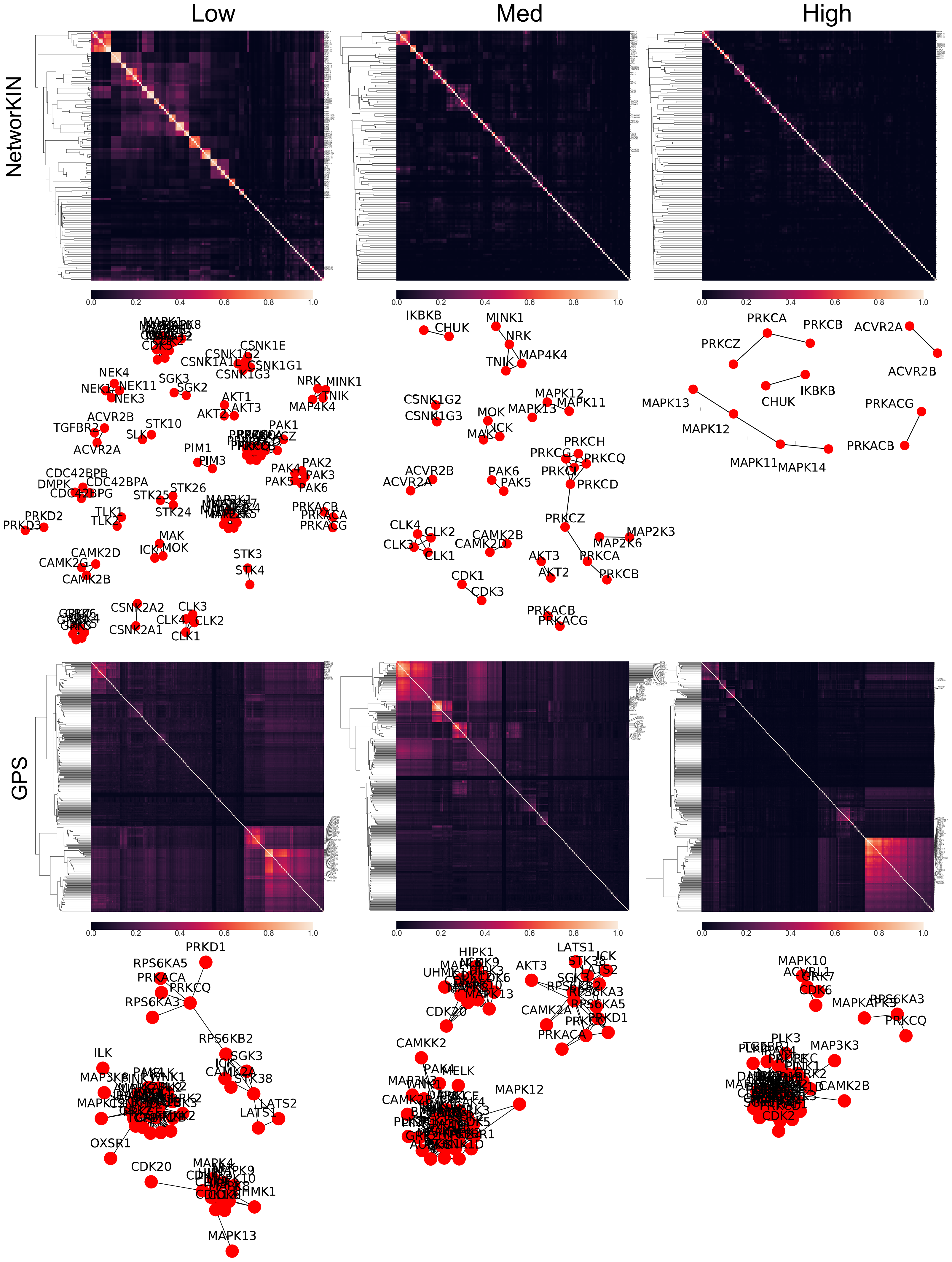

### Supplemental Figure 1: Within Predictor Similarities, continued (ser/thr)

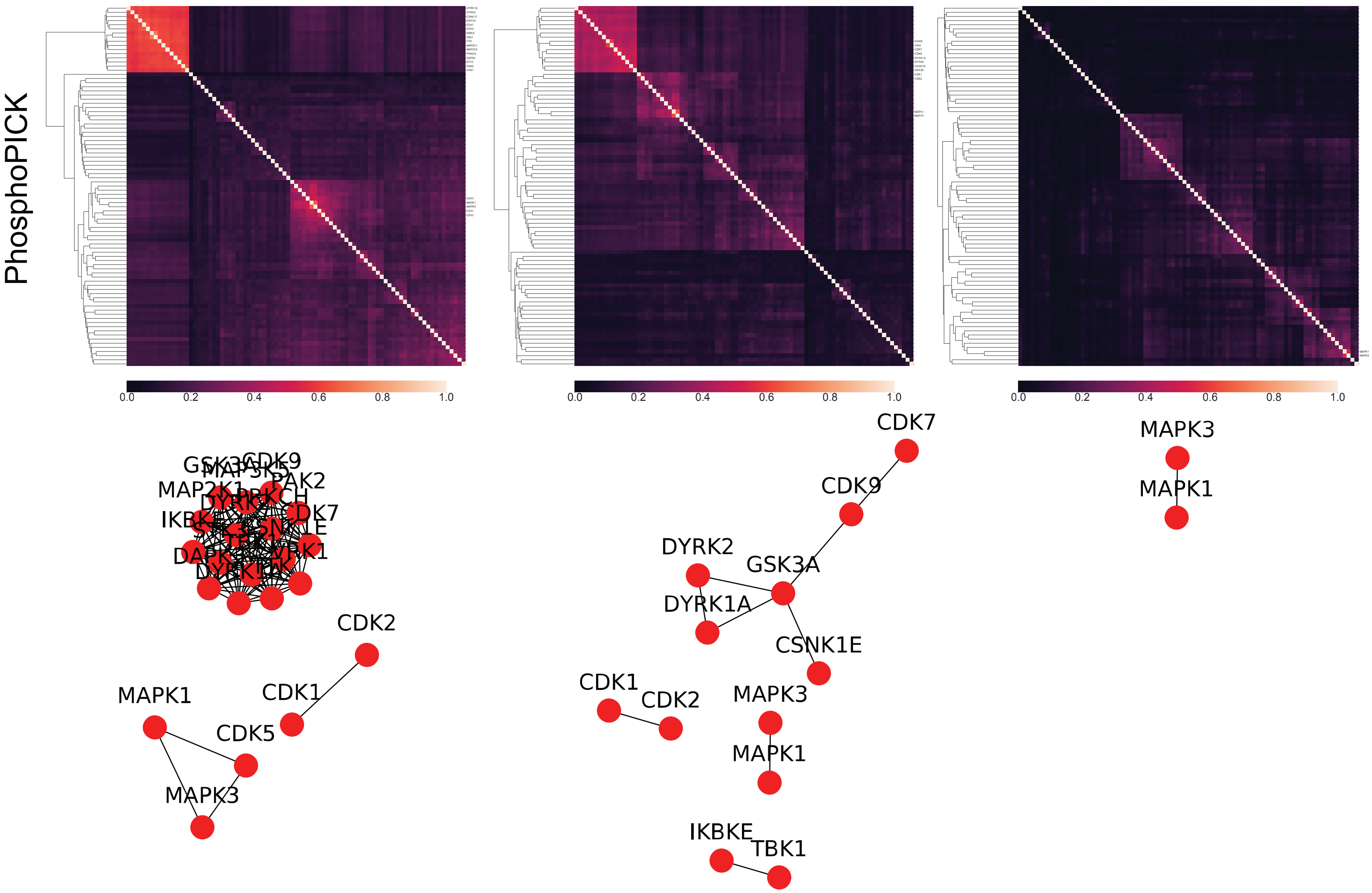

### Supplemental Figure 2: Between Predictor Comparions

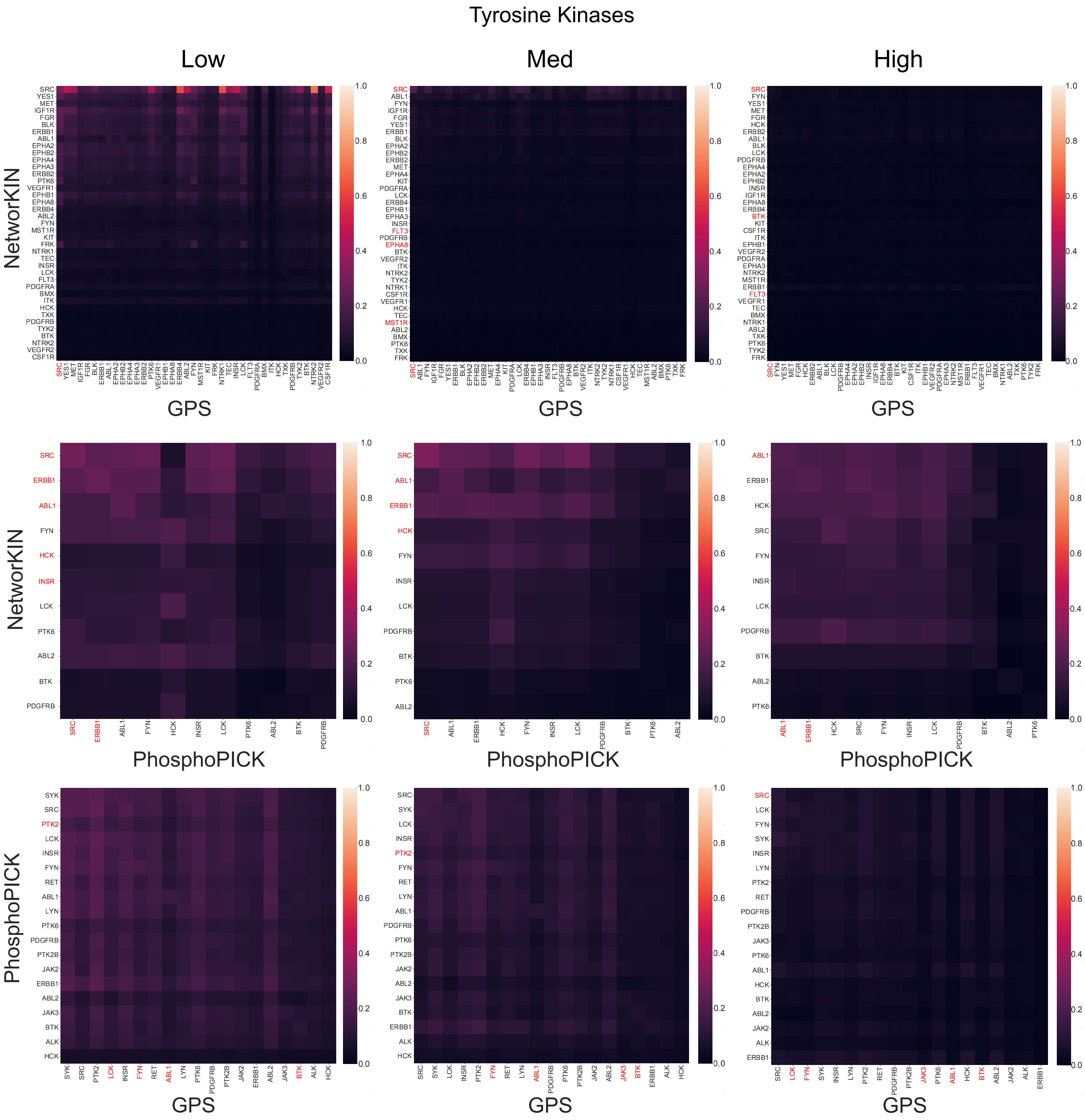

### Supplemental Figure 2: Between Predictor Comparions, continued

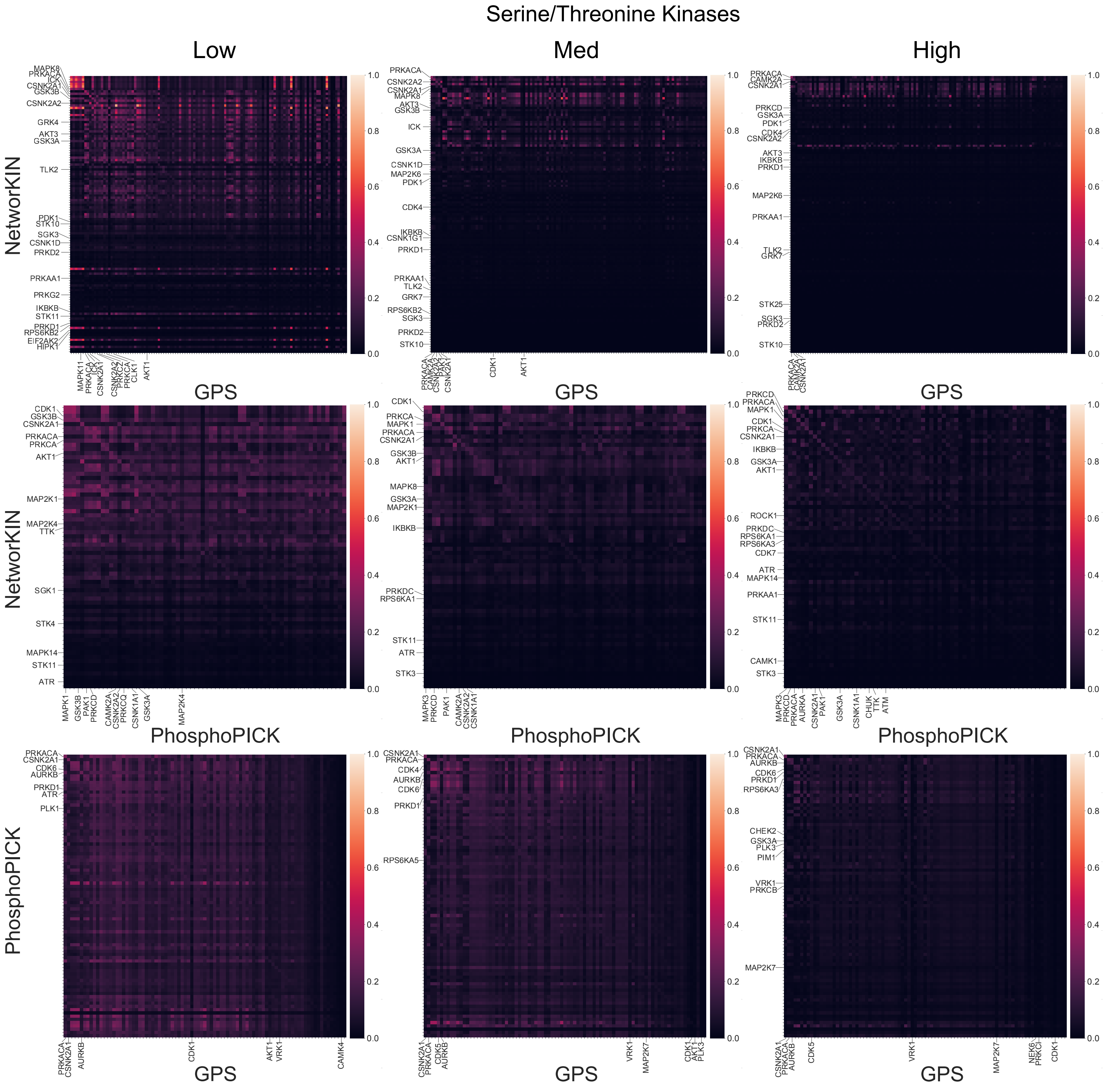

#### Supplemental Figure 3: Ranking Performance for Randomized Datasets

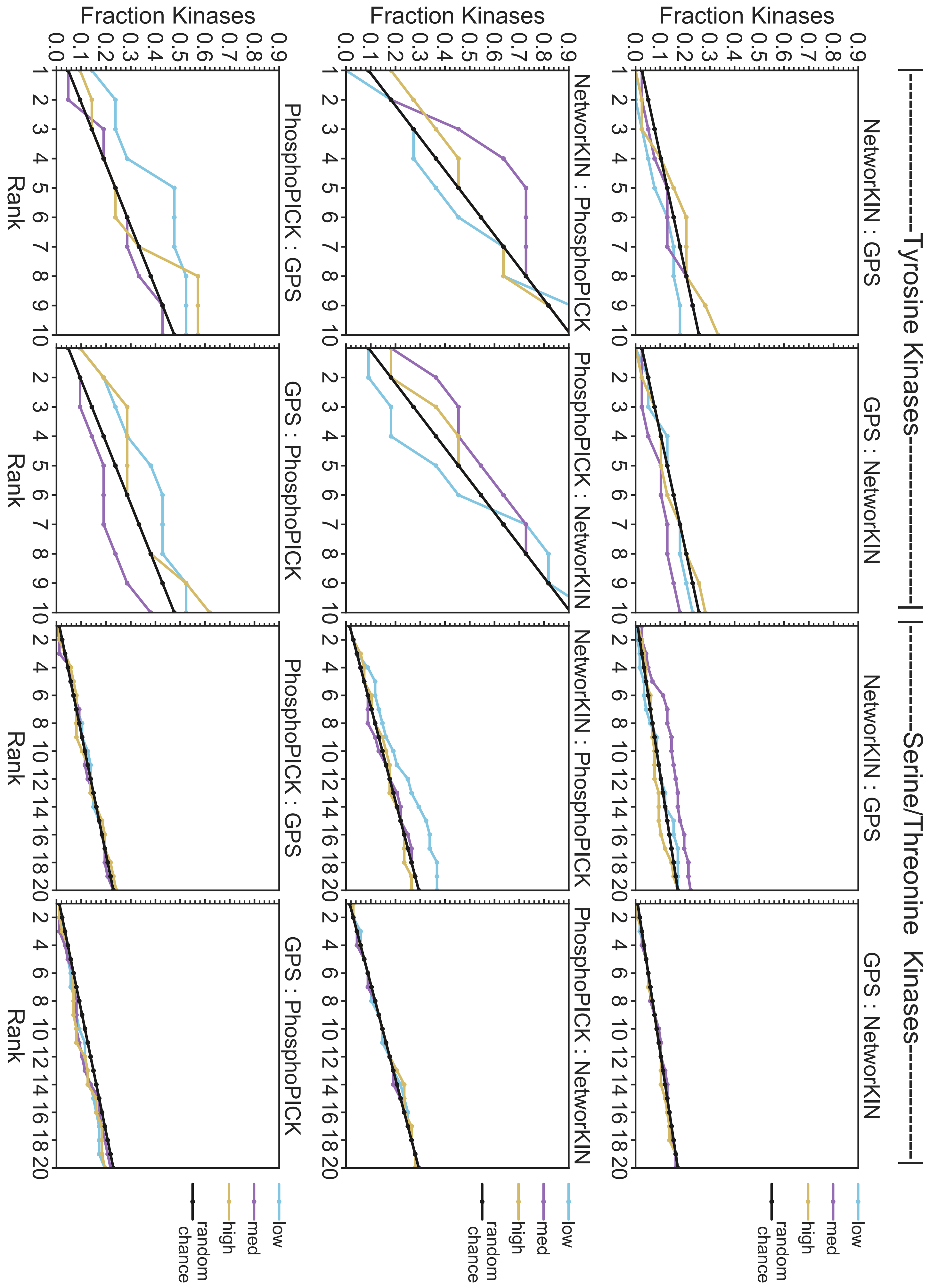
